# Antimicrobial peptide induced-stress renders *Staphylococcus aureus* susceptible to toxic nucleoside analogues

**DOI:** 10.1101/2020.03.30.015206

**Authors:** Alexandro Rodríguez-Rojas, Arpita Nath, Baydaa El Shazely, Greta Santi, Joshua Jay Kim, Christoph Weise, Benno Kuropka, Jens Rolff

## Abstract

Cationic antimicrobial peptides (AMPs) are active immune effectors of multicellular organisms and also considered as new antimicrobial drug candidates. One of the problems encountered when developing AMPs as drugs is the difficulty to reach sufficient killing concentrations under physiological conditions. Here, using pexiganan, a cationic peptide derived from a host defence peptide of the African clawed frog and the first AMP developed into an antibacterial drug, we studied if sub-lethal effects of AMPs can be harnessed to devise treatment combinations. We studied the pexiganan stress response of *Staphylococcus aureus* at sub-lethal concentrations using quantitative proteomics. Several proteins involved in nucleotide metabolism were elevated, suggesting a metabolic demand. We then show that *S. aureus* is highly susceptible to antimetabolite nucleoside analogues when exposed to pexiganan, even at sub-inhibitory concentrations. These findings could be used to enhance pexiganan potency while decreasing the risk of resistance emergence, and our findings can likely be extended to other antimicrobial peptides.

## Introduction

Antimicrobial peptides (AMPs, we use AMPs here as synonymous with host defence peptides) are immune effector molecules used by multicellular organisms to control infections (Nicolas and Mor, 1995; Zasloff, 2002). These peptides are usually active against a broad spectrum of bacterial pathogens and some display activity against antibiotic-resistant bacteria. Thus, antimicrobial peptides are considered a promising source of new antibacterial drugs (Hancock and Sahl, 2006; Czaplewski et al., 2016) to tackle the current antibiotic crisis (Baker, 2015).

Reasons that make AMPs attractive are their high diversity across the tree of life (Wang and Wang, 2004) and the finding that albeit drug resistance evolves also against AMPs (Perron et al., 2006; Habets and Brockhurst, 2012; Lofton et al., 2013; Johnston et al., 2016; Makarova et al., 2018), it evolves at a much lower probability in comparison to conventional antibiotics (Yu et al., 2018; Spohn et al., 2019). One common problem with the development of AMPs as drugs is that, under physiological conditions, their antimicrobial activity cannot easily be recaptured and the required dosage is extremely high (Mookherjee et al., 2020). This dosage issue can be addressed by making use of synergistic combinations of AMPs (Yu et al., 2016), a property common in natural defence cocktails (Westerhoff et al., 1989; Yan and Hancock, 2001).

While the mode of action on bacterial membranes has been worked out for some AMPs (Zerweck et al 2015), the consequence of AMP-induced stress on bacterial physiology is less studied. The first goal of this study, therefore, is to understand how the pathogen *Staphylococcus aureus* reacts to different doses of a pexiganan at the minimum inhibitory concentration (MIC). Pexiganan is a drug that was mostly developed against this bacterium (Ge et al., 1999). This molecule is a 22-amino-acid peptide, an analogue of the magainin peptides isolated from the skin of the African clawed frog. Pexiganan exhibited broad-spectrum antibacterial activity *in vitro* when tested against 3,109 clinical isolates of gram-positive and gram-negative, anaerobic and aerobic bacteria (Ge et al., 1999).

Using pexiganan as an example, we found that different concentrations induce the upregulation of several genes depending on nucleotides or related to nucleotide metabolism. Based on these results, we hypothesized that this will lead to a way to identify phenotypic collateral sensitivity. We hypothesised that the response to pexiganan sensitizes *S. aureus* against certain nucleoside antimetabolites or toxic nucleoside analogues. Interestingly, these analogues have been proposed as an alternative to antibiotics as a consequence of resistance emergence (Thomson and Lamont, 2019). Nucleoside analogues have the advantage of being clinically approved for cancer therapies, but also as antiviral and antifungal treatments (Thomson and Lamont, 2019). Pyrimidine and purine analogues, as we use here, showed potent antimicrobial activity against *S. aureus* in the past (ROGERS and PERKINS, 1960; Stickgold and Neuhaus, 1967; Jordheim et al., 2012a; Thomson and Lamont, 2019).

In this study, we show how proteomic changes of *S. aureus* in response to low-dose pexiganan uncover cellular soft spots that help to identify intervention opportunities. In addition, our findings contribute to the understanding of the early stages of resistance evolution to antimicrobial peptides. Here, we first study the global proteomic response of *S. aureus* to the cationic antimicrobial peptide pexiganan at concentrations similar to and below MIC to detect the possible metabolic changes that open the path to collateral sensitivity to nucleoside analogues. We then confirm that these treatments sensitize *S. aureus* to antimetabolite purine and pyrimidines analogues.

## Material and methods

### Bacteria and growth conditions

We used *S. aureus* SH1000 (Horsburgh et al., 2002) for all experiments. Bacteria were cultured in non-cation-adjusted (unsupplemented) Mueller–Hinton broth (MHB) as recommended for antimicrobial peptides susceptibility testing (Giacometti et al., 2000).

### Global proteomics by LC-mass spectrometry

*Staphylococcus aureus* strain SH1000 was grown in non-cation-adjusted MHB to the mid-exponential-phase (OD_600_ 0.5) at 37°C with vigorous shaking. The cultures were diluted 100 times in fresh MHB in a separate tube to a final volume of 5 ml. Pexiganan was added to tubes for a final concentration of 0.5, 1, 2 and 4 µg/ml (1/8,1/4, 1/2, 1x MIC respectively) in a final culture volume of 10 ml per tube. Non-treated samples were used as controls. After the addition of pexiganan, all tubes were incubated for 30 minutes with moderate shaking at 37°C. The pellets were collected by centrifugation at 10,000 x g for 5 minutes and the supernatant was removed by aspiration using a sterile vacuum line. 50 µl of denaturation urea buffer (6 M urea/2 M thiourea/10 mM HEPES, pH 8.0) were then added to each pellet. The resulting suspensions were transferred to new 1.5 ml Eppendorf tubes and exposed to 5 freeze-thawing cycles alternating between liquid nitrogen and a 37°C water bath. The tubes were centrifuged at 20,000 x g for 10 minutes and the resulting supernatants were transferred to fresh tubes and used as starting protein material for digestion. Each experimental condition had six independent biological replicates. Approximately 50 µg proteins were processed per sample and were in-solution digested as described elsewhere (Rappsilber et al., 2007). Denaturation buffer-containing protein solutions were reduced by adding 1 µl of 10 mM DTT (final concentration) and incubated for 30 minutes. The reactions were then alkylated by adding 1 µl of 55 mM iodoacetamide and incubated for 20 minutes in the dark. Lysyl endopeptidase (LysC, Wako, Japan) resuspended in 50 mM ABC was added to digestion reaction in a proportion of 1 µg per 50 µg of total sample protein and incubated for 3 hours. The samples were diluted with four volumes of 50 mM ammonium bicarbonate (ABC) and digested overnight with 1µg of sequencing grade modified trypsin (Promega, USA). All digestion steps were performed at room temperature. Next day, the digestions were stopped by adding final concentrations of 5% acetonitrile and 0.3% trifluoroacetic acid (TFA). The samples were desalted using the Stage-tip protocol as described previously (Rappsilber et al., 2007), and the eluates were vacuum-dried. Peptides were reconstituted in 10 µl of 0.05% TFA, 2% acetonitrile, and 6.4 µl were analysed by a reversed-phase capillary nano liquid chromatography system (Ultimate 3000, Thermo Scientific) connected to an Orbitrap Velos mass spectrometer (Thermo Scientific). Samples were injected and concentrated on a trap column (PepMap100 C18, 3 µm, 100 Å, 75 µm i.d. x 2 cm, Thermo Scientific) equilibrated with 0.05% TFA, 2% acetonitrile in water. After switching the trap column inline, LC separations were performed on a capillary column (Acclaim PepMap100 C18, 2 µm, 100 Å, 75 µm i.d. x 25 cm, Thermo Scientific) at an eluent flow rate of 300 nl/min. Mobile phase A contained 0.1 % formic acid in water, and mobile phase B contained 0.1% formic acid in acetonitrile. The column was pre-equilibrated with 3 % mobile phase B followed by an increase of 3–50% mobile phase B in 50 min. Mass spectra were acquired in a data-dependent mode utilising a single MS survey scan (m/z 350–1500) with a resolution of 60,000 in the Orbitrap, and MS/MS scans of the 20 most intense precursor ions in the linear trap quadrupole. The dynamic exclusion time was set to 60 s and automatic gain control was set to 1×10^6^ and 5,000 for Orbitrap-MS and LTQ-MS/MS scans, respectively.

MS and MS/MS raw data were analysed using the MaxQuant software package (version 1.6.4.0) with implemented Andromeda peptide search engine (Tyanova et al., 2016a). Data were searched against the reference proteome of *S. aureus* strain NCTC 8352 downloaded from Uniprot (2,889 proteins, taxonomy 93061, last modified September 2017) using label-free quantification and match between runs option was enabled. Filtering and statistical analysis was carried out using the software Perseus (Tyanova et al., 2016b). Only proteins with intensity values from at least 3 out of 6 replicates were used for downstream analysis. Missing values were replaced from normal distribution (imputation) using the default settings (width 0.3, down shift 1.8). Student’s T-tests were performed using permutation-based FDR of 0.05.

### Antimetabolite nucleosides

In this study, we used four nucleoside analogues. We used the pyrimidine analogues 6-azauracil, gemcitabine, 5-fluorouracil and the purine analogue 6-thioguanine. All drugs were purchased from Sigma Aldrich (Germany). 6-azauracil is used as a growth inhibitor of various microorganisms via depletion of intracellular GTP and UTP nucleotide pools (Habermann, 1961). Gemcitabine is a chemotherapy medication used to treat different types of cancer. Gemcitabine is a synthetic pyrimidine nucleoside analogue in which the hydrogen atoms on the 2’ carbon of deoxycytidine are replaced by fluorine atoms and competitively takes part and disrupts several pathways where pyrimidines are needed (Jordheim et al., 2012b). 5-Fluorouracil is also used as an anticancer treatment and it works by inhibiting cell metabolism by blocking many pathways, but its major action is the inhibition of the thymidylate synthase. By doing so, the synthesis of the pyrimidine thymidine is stalled, which is an essential nucleoside required for DNA replication (Singh et al., 2015). 5-Fluorouracil causes a drop on dTMP and causing cells to undergo cell death via thymineless death (Khodursky et al., 2015; Singh et al., 2015).

### Pexiganan and antimetabolite nucleosides susceptibility testing

Minimal inhibitory concentration (MIC) was determined by broth micro-dilution method modified for cationic antimicrobial peptides (Wiegand et al., 2008). Briefly, 2 µl of the mid-exponential phase culture diluted 1:100 (around 10^5^ bacteria) were inoculated into each well of a polypropylene V-bottom 96-well plates with anti-evaporation ring lids (Greiner Bio-One GmbH, Germany). Prior to inoculation, pexiganan and the analogues (a kind gift from Dr Michael A. Zasloff, Georgetown University) were two-fold serially diluted in a final volume of 100 µl MHB per well using 32 µg/ml as starting concentration. Each assay was performed with eight replications and plates were incubated at 37°C in a humid chamber. The MIC was defined as the lowest concentration where no visible bacterial growth was observed after 24 hours.

### Isobologram Assay

The combined activity and interactions between peptides and pexiganan and purine and pyrimidine analogues against *S. aureus* in MHB was determined using isobolographic combinations, also called checkerboard assay method, (8×8 matrix of concentrations combinations) (Tallarida, 2006). In a 96-well plate, 50 μl of pexiganan at 4x MIC concentration was two-fold serially diluted ranging from 32 to 0.25 μg/ml in the direction of the columns from 1 to 8. In another 96-well plate 100 μl of nucleoside analogues at 8x MIC concentrations were prepared in an identical way as the previous plate, but diluted in the direction of the rows from A to H. Half of the content (50 μl) of each well from the analogue drug plate was transferred to the corresponding well of the plate containing pexiganan in an equal 1:1 mix fashion, halving the concentration of both compounds. In the same plate, the columns 9 and 10 were used to serially dilute both, the peptide and the analogue drug in the same concentrations that were present in the combination to compare single compounds vs combination. Columns 11 and 12 were used as a control, by inoculating column 11 wells with bacteria without any drug and leaving columns 12 only with the same volume of MHB as a media contamination control. Each plate was prepared in triplicates to check for consistency. The bacterial suspension was prepared by growing *S. aureus* SH1000 to mid-exponential phase (2.5 hours, with moderate shaking at 37°C) in MHB to an OD_600_ between 0.3 to 0.5. The bacterial suspension was diluted in MHB and approximately 1 × 10^6^ bacteria were inoculated in each well. After 24 hours of incubation at 37°C in a humid chamber, the plates were visually examined for growth. The Fractional inhibitory concentration (FIC index) for a combination of pexiganan and each antimetabolite drug was calculated as [(MIC of the peptide in combination with a given analogue)/(MIC of peptide alone)] + [(MIC of analogue in combination with peptide)/(MIC of analogue alone)]. The interpretation of the results was as follow: FIC ≤ 0.5, synergistic; 0.5 < FIC ≤ 1, additive; 1 < FIC ≤ 4, indifferent; FIC > 4, antagonistic, antagonistic (Ng et al., 2018). To ensure that bacteria lost viability while reading MIC values for pexiganan-analogue combinations, we used the resazurin colorimetric assay as described previously with minor modifications (Elshikh et al., 2016). Resazurin (THK, Germany) was prepared at 0.015 % in distilled water and sterilised by filtration. It was stored at 4°C for a maximum of 1 week after preparation. Resazurin (0.015 %) was added to each well (10 µl per well, 1/3 of the original described quantity) and further incubated for 3 hours for the observation of colour change. Columns with no colour change (blue resazurin) were scored as dead culture. In contrast, colour change to purple (reduced resazurin) was considered as a sign of viability.

### Time-kill experiments

Starting from early mid-exponential phase cultures (1×10^7^ CFU/ml), bacteria were exposed to growing concentrations of pexiganan ranging from 1 to 8x MIC or pexiganan combined with the nucleoside analogues 6-azauracil, gemcitabine, 5-fluorouracil and 6-thioguanine at their respective 1/2x MIC values. The cultures were incubated with soft shaking at 37°C for 2 hours. Samples from each culture (1ml) were taken at 20-minute time-point intervals. The samples were diluted and plated to determine cell viability. The experiments consisted of five independent replicates. Non treated cells were used as a control.

### Statistical analysis

The effect of treatments on bacterial killing was analysed using R package nparLD (Noguchi et al., 2012). *P* values less than or equal to 0.05, after correction, if needed, were considered statistically significant. All tests were performed with the statistic software R (R Core Team, 2017).

## Results

### Changes in protein profiles after pexiganan treatment

We examined *S. aureus* exposed to pexiganan by studying proteome-wide changes after a 30-minute treatment with different pexiganan concentrations (0.125, 0.25, 0.5 and 1x MIC, Table S1). Overall, 1160 proteins were identified at 1 % or less false discovery rate (FDR) among which 968 proteins were quantified in at least 3 out of 6 replicates and used for downstream analysis. All identified proteins, their quantification and statistical tests are provided in supplementary Table S2. A global overview shows a proteome-wide perturbation induced by pexiganan stress for all concentrations compared to control. Many proteins were significantly differentially expressed, Figure S1). It is noticeable that as long as the dose increases, the level of expression (fold-change) of both overexpressed and suppressed genes, decreases, making the dot scattering of the volcano plot less disperse (Figure S1). This indicates a decrease in the ability of the cell to react with increasing peptide concentration.

Within the upregulated proteome fraction (Figure 1, Figure S1), a group of proteins related to osmotic stress response shows up. The multi-peptide resistance factor MprF, a protein associated with cationic peptide resistance, which is conserved among many bacterial species (Kristian et al., 2003; Weidenmaier et al., 2005) is upregulated in all pexiganan doses except in the lower one (1/8x MIC). MprF catalyses the transfer of a lysyl group from L-lysyl-tRNA(Lys) to membrane-bound phosphatidylglycerol producing lysyl-phosphatidylglycerol, a major component of the bacterial membrane with a net positive charge, hence modification of anionic phosphatidylglycerol with positively charged L-lysine results in the repulsion of the peptides. Changes of the membrane charge is a *per se* resistance mechanism against cationic antimicrobial peptides. Thus, MprF increases resistance to moenomycin and vancomycin, resistance to human defensins (HNP1-3) and evasion of oxygen-independent neutrophil killing and other AMPs and antibiotics (Oku et al., 2004; Staubitz et al., 2004). Another highly expressed protein is CapFis, involved in the pathway capsule polysaccharide biosynthesis, a mucous layer on the surface of the bacterium that facilitates immune evasion and infection. CapF is an important virulence factor during infections by *S. aureus*. The enzyme CapF is considered a therapeutic candidate to disrupt the capsule polysaccharide biosynthesis (Miyafusa et al., 2013). TagG upregulates the protein which is part of the wall teichoic acid synthesis during the final steps of the pathway. Wall teichoic acids are important in pathogenesis and play key roles in antimicrobial resistance (Weidenmaier et al., 2005; Brown et al., 2013). The chaperons/proteases ClpL and TreP are among the fifty upregulated genes for the dose corresponding to the MIC (8 μg/ml). ClpL is an ATP-dependent Clp protease. Clp proteases play a central role in stress survival, virulence and antibiotic resistance of *S. aureus* (Frees et al., 2014).

**Figure 1.**
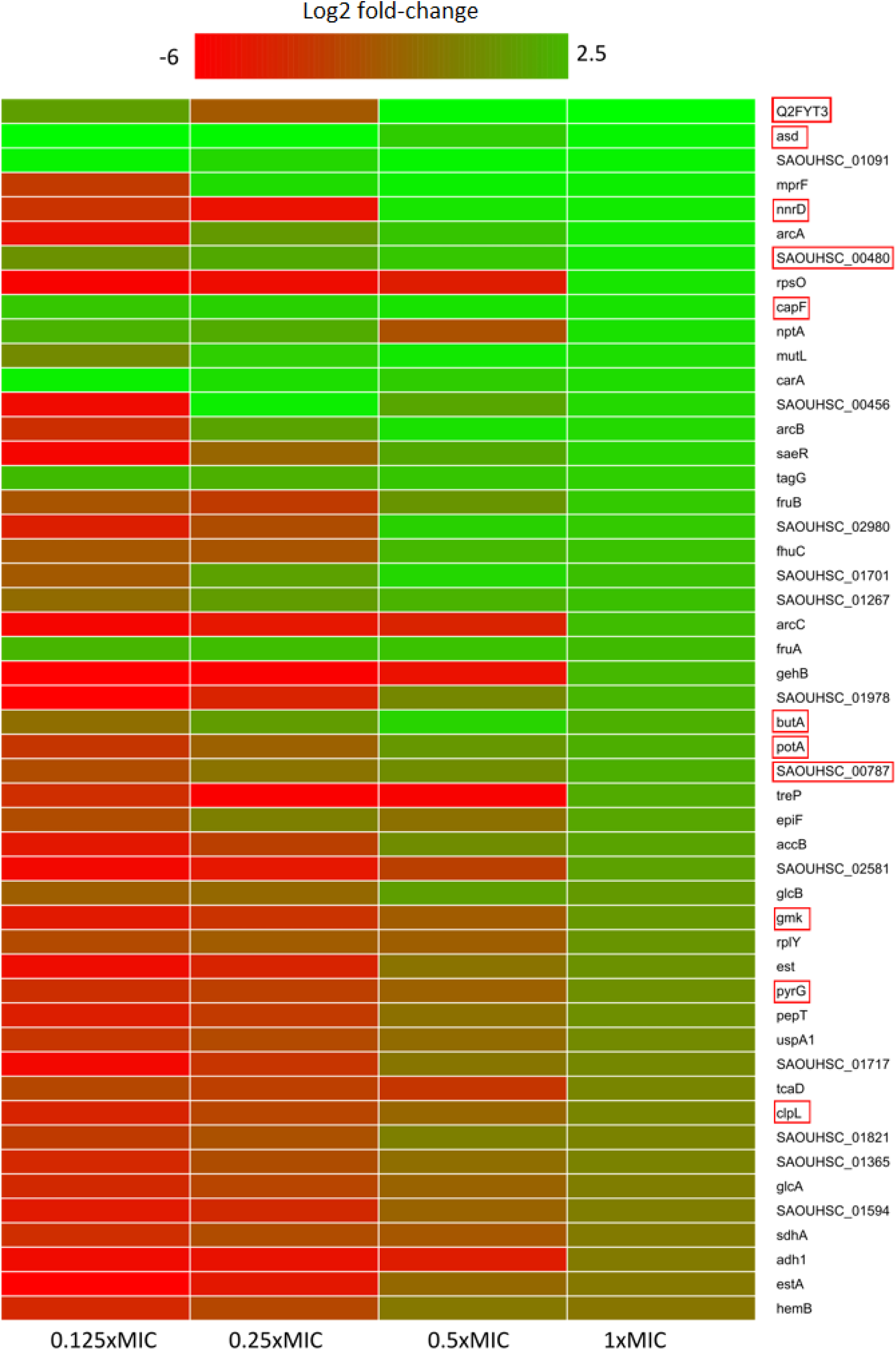
Heatmap of relative protein expression based on label-free quantification detected by liquid chromatography-mass spectrometry (LC-MS). Only the 50 most significantly up-regulated proteins compared to the control at 1XMIC are shown (log2 fold-change). Red rectangle highlights proteins that participate in or depend on nucleotide metabolism. Proteins were extracted after 30 minutes of pexiganan addition (0.125, 0.25, 0.5 and 1 fractions of the minimal inhibitory concentration). Intensity ranges of the log2 fold-changes are given from highest intensity (green) to lowest (red).

In addition to the above virulence factors, pexiganan also induced the expression PepT, also known as Staphopain A. This enzyme is a cysteine protease that plays an important role in the inhibition of host innate immune response. It cleaves host elastins from connective tissues, pulmonary surfactant protein A in the lungs, and the chemokine receptor CXCR2 on leukocytes (Kantyka et al., 2013). Proteolytic cleavage of surfactant protein A impairs bacterial phagocytosis by neutrophils while CXCR2 degradation blocks neutrophil activation and chemotaxis (Potempa et al., 1988; Kantyka et al., 2013). Additionally, PepT promotes vascular leakage by activating the plasma kallikerin/kinin system, resulting in patient hypotension (Imamura et al., 2005). Another important virulence factor is coded by the Gene SAOUHSC_02980, a protein containing an isochorismatase domain. This enzyme participates in the biosynthesis of siderophore groups that in *S. aureus* are redundant systems and varies even across different vertebrates hosts (Perry et al., 2019).

NptA, a phosphate transporter, usually induced by phosphate limitation, is highly abundant. Inorganic phosphate acquisition via NptA is particularly important for the pathogenesis of *S. aureus*. NptA homologs are widely distributed among bacteria and closely related less pathogenic staphylococcal species do not possess this importer. Another phosphate metabolism-related gene with high expression is SAOUHSC_00480, that codes for a putative nucleoside triphosphate pyrophosphohydrolase (Pundir et al., 2016). Another two proteins, CarA and CarB, participate in the L-arginine biosynthesis. They are involved in the first step of the sub-pathway that synthesizes carbamoyl phosphate from bicarbonate. The elevation of these enzymes could indicate that pexiganan stress may be involved in amino acid depletion. Also related to phosphate metabolism, we observed a high level of FruA in different pexiganan concentrations. This protein a phosphoenolpyruvate-dependent sugar phosphotransferase system (a PTS system) is a major carbohydrate active transport system, which catalyses the phosphorylation of incoming sugar substrates concomitantly with their translocation across the cell membrane and potentially important for survival in the respiratory tract of the host (Garnett et al., 2014). GlcB, another PTS system is a phosphoenolpyruvate-dependent sugar phosphotransferase system. This protein is another major carbohydrate active -transport system and catalyses the phosphorylation of incoming sugar substrates and their translocation across the cell membrane (Vitko et al., 2016).

The gene SAOUHSC_00456 that codes for YabA is significantly increased as well. YabA is involved in the initiation of chromosome replication and is a negative controller of DNA replication initiation in *Bacillus subtillis*. YabA and DnaD inhibit helix assembly of the DNA replication initiation protein DnaA (Scholefield and Murray, 2013). The elevated concentration within the cell of YabA could stall the cell division while the bacteria is under severe stress. *S. aureus* upregulates Spermidine/putrescine import ATP-binding protein PotA. This protein is part of the ABC transporter complex PotABCD and responsible for energy coupling to the transport system. Spermidine and putrescine are polyamines which role in *S. aureus* is not well defined (Di Martino et al., 2013). There are also a set of up-regulated proteins coded by the genes SAOUHSC_01717, SAOUHSC_02581 and SAOUHSC_02581 which function remains unknown as described in Uniprot database and showed no homology with any known sequence (Pundir et al., 2016).

One of the hallmarks of our proteomic dataset is that we found a higher level of expression, compared to control, for proteins related with nucleotide metabolism (Figure 1), which is directly connected to the upregulation of phosphate metabolism proteins described above. GmK for example, an essential protein for recycling GMP and indirectly, cGMP Guanylate kinase is highly upregulated. GMK is an essential enzyme and a potential antimicrobial drug target owing to its role in supplying DNA and RNA precursors (Omari et al., 2006). Another nucleobase metabolism-related protein having or exhibiting a higher expression for the 1x MIC treated cells is PyrG. This enzyme catalyses the ATP-dependent amination of UTP to CTP with either L-glutamine or ammonia as the source of nitrogen. It also regulates intracellular CTP levels through interactions with the four ribonucleotide triphosphates.

Pexiganan also negatively impacted the level of expression of many proteins, proteome-wide (Figure S1, Table S2). Among the most affected gene expressions throughout all concentrations are genes such as *dps* (coding for a known iron storage protein), *hld, copZ, cspC, metQ, sceD, isaA csoB*/*C, dltC*, adsA and *sasG*, SAOUHSCA_01134 and SAOUHSCA_02576. The gene cspB codes for the downregulated protein CspD, a cold shock protein that accumulates during low temperature or cold shock. This gene is also a component of the stringent response, indicating that it could be a general stress response gene (Anderson et al., 2006). Other genes showing a differentially low level of expression are SAOUHSC_01986, SAOUHSC_01986, SAOUHSC_008020, SAOUHSC_02093, SAOUHSC_02535 and SAOUHSC_01414 which code for uncharacterized proteins (Pundir et al., 2016). SAOUHSC_01030 is a putative glutaredoxin domain-containing protein but it is not characterized either. The gene SAOUHSC_02576 codes for a putative secretory antigen SsaA, identified in *S. epidermidis* but its function is also unknown (Pundir et al., 2016).

In contrast to the upregulation of peptidoglycan synthesis, we observe that putative peptidoglycan hydrolases and probable lytic transglycosylases IsaA and SceD were downregulated. Interestingly, the *isaA sceD* double mutant is attenuated for virulence, while SceD is essential for nasal colonization in cotton rats (Stapleton et al., 2007). The gene *moaD* shows also a reduced level of expression and it codes for a molybdopterin converting factor subunit 1. Molybdopterins are a class of cofactors found in most molybdenum-containing and all tungsten-containing enzymes. Molybdopterin pathway reactions consume guanosine triphosphate that is converted into the cyclic phosphate of pyranopterin (Mendel and Leimkühler, 2015). Another metabolic enzyme, AldA, aldehyde dehydrogenase central carbohydrate metabolism is downregulated in all doses of pexiganan. This is also the case of CopZ, a chaperone that serves for the intracellular sequestration and transport of copper, delivering it to the copper-exporting P-type ATPase A (CopA) (Sitthisak et al., 2007).

### Pexiganan stress has a strong impact on the essential proteome

We visualised the global impact of pexiganan stress (at 1x MIC) on bacterial physiology by a network analysis based on protein-protein interactions and function (Szklarczyk et al., 2015) of *S. aureus* essential genes (Figure S3). This network analysis provides global view information on protein level alterations and integrates protein-protein interactions, including indirect functional and direct physical associations (Szklarczyk et al., 2015). At this concentration, it is noticeable that the majority of the essential genes are downregulated, and it is possible that this pattern has strong influence on pexiganan lethality. The majority of upregulated proteins are ribosomal components.

### Gene ontology analysis points to an upregulation of nucleotide metabolism

The signature of pexiganan stress on *S. aureus* in the upregulated fraction points to nucleotide metabolism-related genes. GO annotation allows enrichment analysis providing global information based on the gene expression levels by proteomics or transcriptomics or other gene expression datasets (Mi et al., 2019). We focus this comparative analysis on the protein expression levels of the most 100 most upregulated proteins of every pexiganan dosage. We focussed on categorizing by pathways. Some of the upregulated pathways involved genes related to oxidative stress, peptidoglycan synthesis and N-acetylglucosamine that are expected from cationic antimicrobial peptides since they attack the cell envelopes. In addition, there is a reactivation of the central metabolism by the upregulation of genes from glycolysis, TCA cycle, arginine and thiamine synthesis. However, the most enriched pathways in the GO analysis for all pexiganan concentrations were related to nucleotide metabolism (Figure 2). The nucleotide upregulated pathways include ATP synthesis, Adenine and hypoxanthine salvage pathways, de novo synthesis of purines and pyrimidines and S-adenosylmethionine. This result confirms that pexiganan stress induces a scarcity of these metabolites within the cell. Taking into account the previous results, we hypothesised that upregulation of nucleotide-dependent and related genes could create a collateral sensitivity.

**Figure 2.**
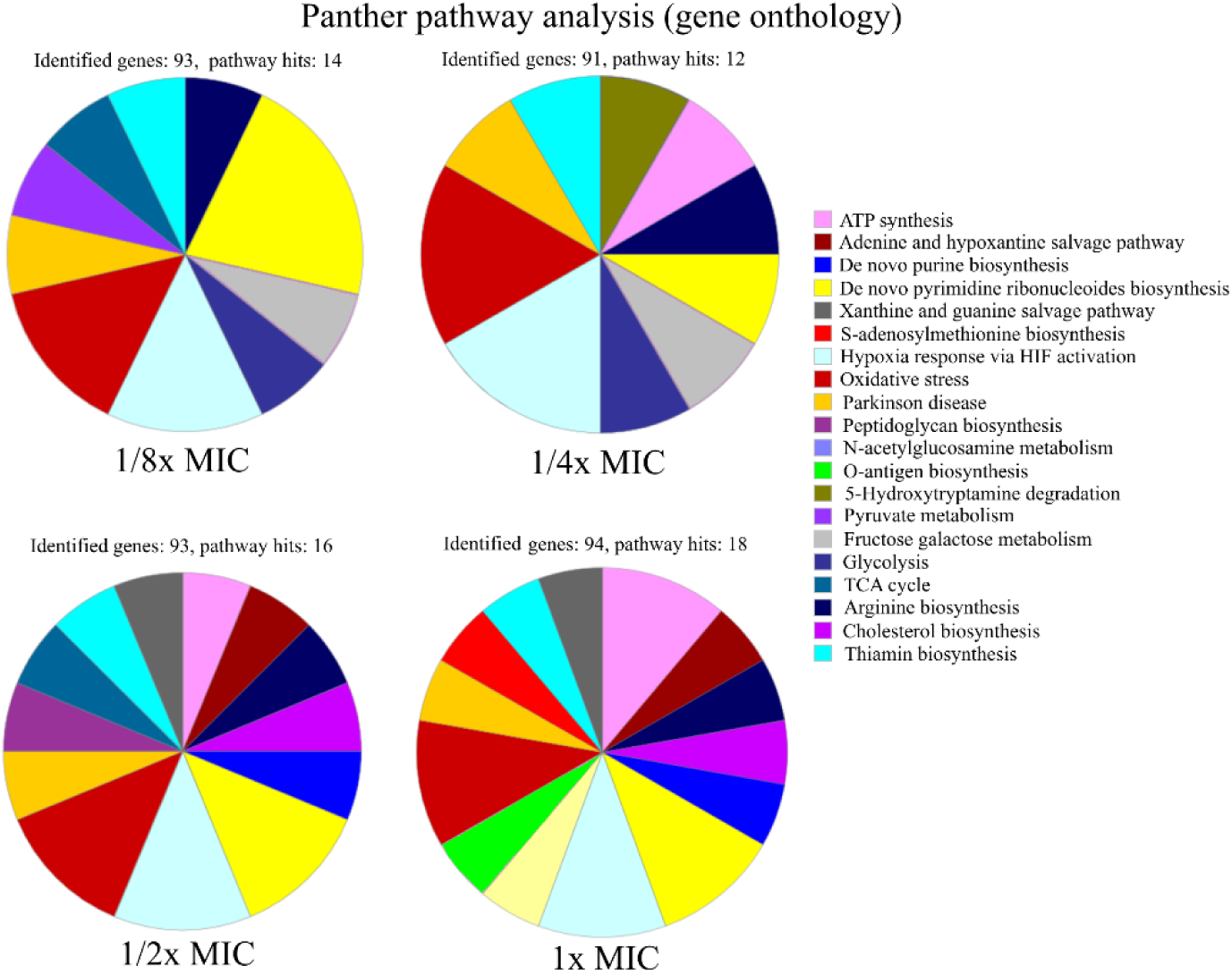
Functional characterisation of pathways of up-regulated proteins in *S. aureus* SH1000 at different concentrations of pexiganan (0.125, 0.25, 0.5 and 1 fractions of the minimal inhibitory concentration). For this analysis, only the 100 most highly differentially expressed proteins for each concentration of pexiganan were used. The analysis was carried out using the online gene ontology analysis software PANTHER (Mi et al., 2019).

### Nucleoside analogues antimetabolites act synergistically with pexiganan

We designed a simple drug interaction experiment between pexiganan and some nucleoside analogues including the purine and pyrimidines antimetabolites: 6-azauracil, gemcitabine, 5-fluorouracil and 6-thioguanine (Figure S4, Figure S5, Table S4). This experiment is the classic isobologram, also known as checkerboard assay (Tallarida, 2006). All analogues showed a synergistic activity when combined with pexiganan (Table S4). The most active ones were 5-fluorouracil and gemcitabine and, while the 6-azauracil and 6-thiogunine showed a milder effect according to their respective Fractional inhibitory concentration index (Table S4). All the combinations managed in all cases a decrease of the minimal inhibitory concentration for each drug when compared to the respective drug alone. These results indicate pexiganan induces a strong collateral sensitivity to nucleoside analogues.

To study the influence of nucleoside analogues on the killing by pexiganan, we carried out a time-kill experiment combining each of 6-azauracil, gemcitabine, 5-fluorouracil and 6-thioguanine with pexiganan. We assayed all drugs using half of the minimal inhibitory concentration. We exposed mid-exponential phase *S. aureus* cells to these combinations and sampled the viability of the cultures every 20 minutes (Figure 3). All compounds significantly increased the killing ability of pexiganan, gemcitabine and 5-fluorouracil being the most active drugs. The killing rate was increased by some order of magnitudes in all combinations. The killing by the combination of gemcitabine or 5-fluorouracil with pexiganan, at their corresponding half MIC values, was more efficient than 8x MIC concentration of pexiganan alone. The viability was assessed not only by the absence of growth but also by the addition of resazurin, a reagent that turns from blue to purple when it is reduced by microbial enzymes that only work within living bacteria (Elshikh et al., 2016).

**Figure 3.**
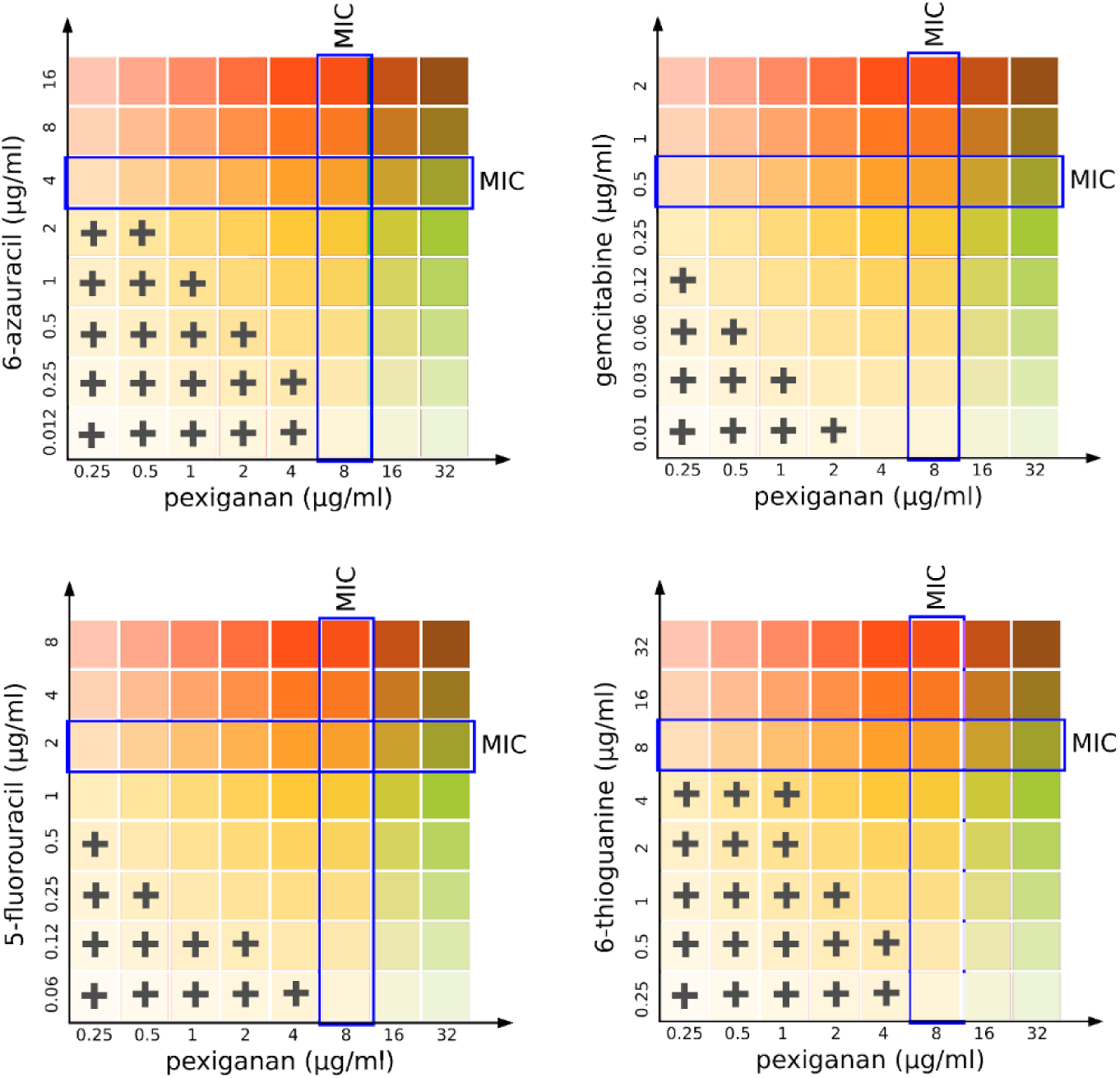
Isobolographical response of pexiganan combination with different antimetabolite nucleosides. Blue squares represent the minimal inhibitory concentrations for pexiganan and each of the tested analogues (see also supplementary Figure S5). The crosses indicate the presence of bacterial growth in the unique concentration combinations of each well.

**Figure 4.**
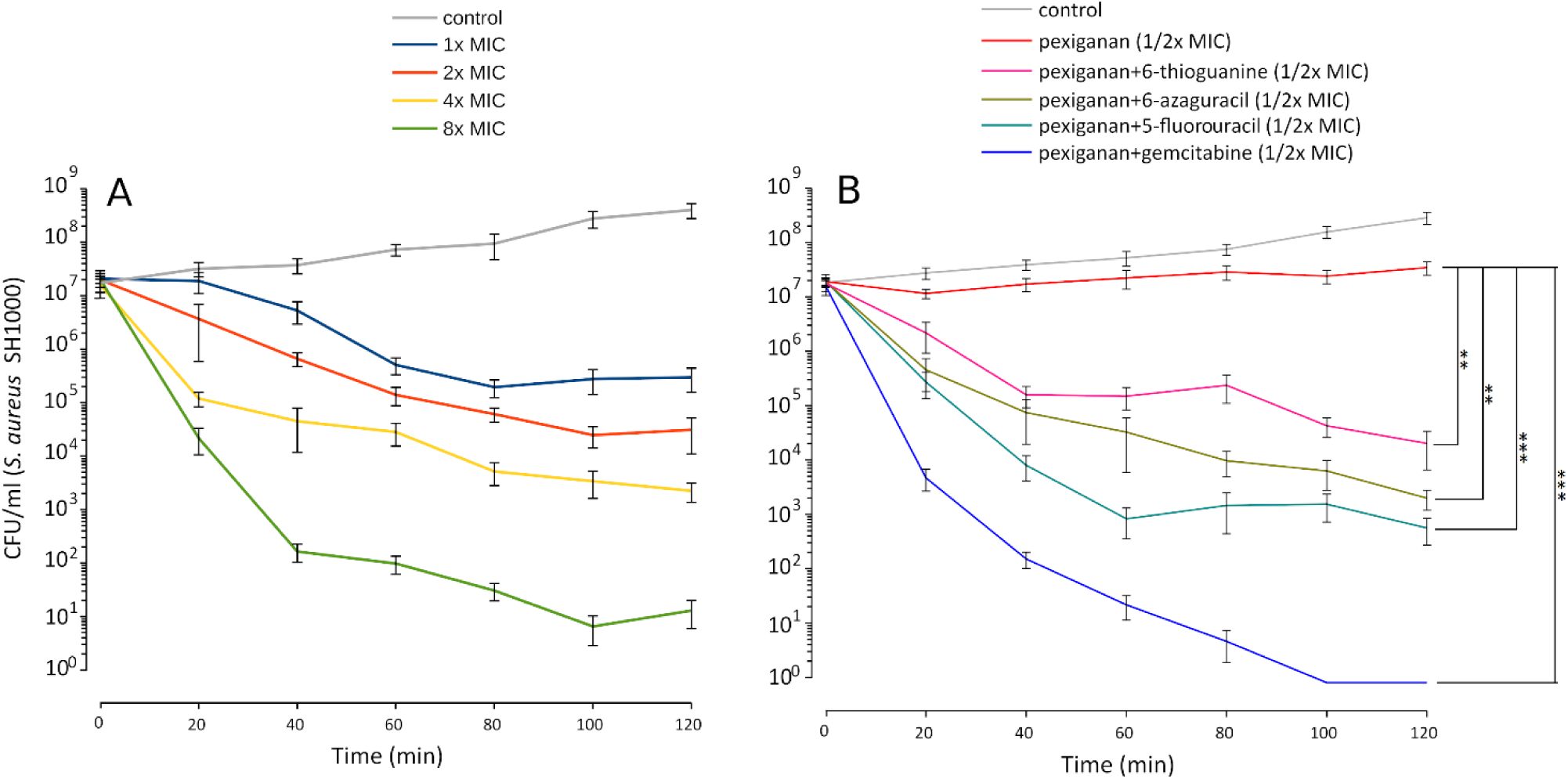
Pexiganan-nucleoside antimetabolite combination drastically increases the killing capacity of pexiganan. (A) Killing dynamic of *S. aureus* SH100 at different concentrations of pexiganan using the MIC as the starting point. (B) Example data of time-kill experiment exposing mid-exponential phase bacterial cultures to pexiganan-nucleoside antimetabolite combinations (both at 1/2x MIC concentrations). The combination has a dramatic effect on the killing ability of pexiganan. Data points were determined by counting colony-forming units (CFU) at different time points. Mean ± SDM, n=5. Asterisks represent significant differences (R package nparLD, one asterisk for p<0.05 and two asterisks for p<0.01 and three asterisks for p<0.001). Only comparisons between pexiganan (1/2x MIC) and pexiganan-analogues combinations are shown.

## Discussion

We have found that pexiganan, a cationic antimicrobial peptide, can induce a stress response in *S. aureus* that results in a proteome-wide impact. Pexiganan treatment upregulates known virulence factors such as MprF, the capsule synthesis protein CapF, a wall teichoic acid TagG, the proteases ClpL and PepT and other proteins important for the interactions with the hosts. This could lead to a phenotypic cross-tolerance of other immune effectors of hosts and possibly complicate the bacterial infection in case of inefficient treatment where bacteria could be exposed to sub-lethal concentrations. This is a legitimate concern since AMP-resistant variants have been reported to have evolved that have shown some cross-resistance with immune system effectors (Bell and Gouyon, 2003; Fleitas and Franco, 2016). This risk has been shown for pexiganan as well (Habets and Brockhurst, 2012). Our data also provides input about possible induced physiological changes that would help *S. aureus* to adapt to the intra-host environment during its interaction with specific immune system effectors.

It is important to note that, given the coverage of the proteomic data and range of pexiganan doses, we did not find evidence of activation of mutagenic stress pathways or recombination. This indicates that the mode of killing by cationic antimicrobial peptide does not increase genome instability as is typical for classic antibiotics (Blázquez et al., 2012). Along these lines, we have previously shown and proposed that antimicrobial peptides, including pexiganan, do not increase neither mutagenesis (Rodríguez-Rojas et al., 2014) nor recombination (Rodríguez-Rojas et al., 2018) in Gram-negative bacteria. Our findings here are consistent with these observations in the Gram-positive model bacterium *S. aureus*.

The elevated level of expression of proteins such as GmK, PyrG, NptA and some amino acids-related enzymes such as CarA and CarB that participate in the biosynthesis of L-arginine could be explained by changes in permeability. Amino acids, nucleobases and nucleotides are small molecules that could easily escape from the cellular compartment in case of membrane damage. This is a well-known property of cationic agents, including AMPs (Asthana et al., 2004; Brogden, 2005; Rodríguez-Rojas et al., 2015). The fact that only a few proteins from the amino acids biosynthesis pathways are upregulated could be explained because the experiments were carried out in a complex medium like MHB that contains several amino acids and bacteria would upregulate only necessary pathways. A similar situation might be expected within a host.

The upregulation of the phosphate and nucleotide-related proteins provides a direction to investigate drug susceptibilities created by pexiganan stress. Although the antimetabolites used in this work have good antibacterial activity, if they are used in monotherapy they are also prone to generate resistance (Jordheim et al., 2012c; Thomson and Lamont, 2019). Thus, their use in combination could possibly help to prevent resistance (Yu et al., 2016; Tyers and Wright, 2019).

The synergistic combined action of pexiganan with nucleoside antimetabolites could be explain probably by two underlying mechanisms. First, pexiganan stress forces a response by *S. aureus* that upregulates nucleobase salvage pathways and other nucleotide-dependent metabolic pathways. Second, pexiganan has the potential to change membrane permeability and induce the uptake of such metabolites even at sublethal concentrations possibly leading to much higher intracellular concentrations (Figure 5). We have shown previously that cationic antimicrobial peptides can mediate the uptake of small molecules due to changes in permeability at sublethal concentrations (Rodríguez-Rojas et al., 2015). The more potent activity of gemcitabine and 5-fluorouracil could be explained because they act on the cell walls as previously reported (Gieringer et al., 1986; Jordheim et al., 2012c).

**Figure 5.**
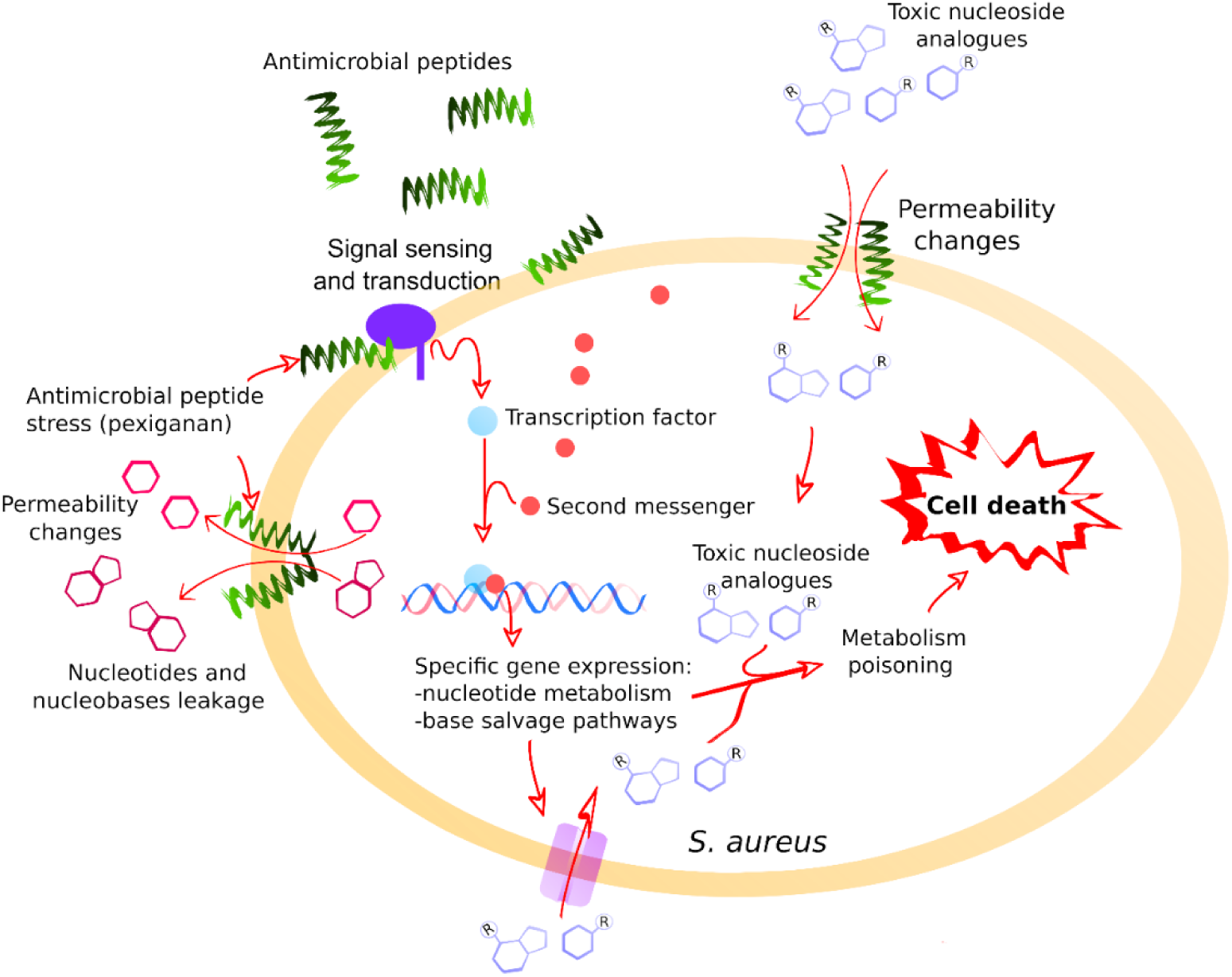
A general model illustrating the positive interaction between pexiganan and nucleoside antimetabolites against *S. aureus*. The interactions of pexiganan with the membrane at sub-inhibitory concentrations lead to transient permeability changes in the envelope that promote leakage of small molecules such as nucleotides, nucleobases or nucleosides. Simultaneously, other small molecules such as toxic nucleoside analogues can increase the diffusion rate toward the intracellular compartment. This stress is sensed by the cell that responds by activating nucleoside metabolism creating an intervention opportunity. In this situation, toxic nucleoside antimetabolites are more efficiently incorporated into RNA, DNA other nucleotide depending reactions that may include envelope synthesis, enhancing toxicity and leading to faster cell killing.

An additional potential therapeutic advantage of the nucleoside analogues studied here is that all the clinical properties of these drugs are well known, including toxicological profile, pharmacological activities and metabolising properties (Cheng et al., 2018; Thomson and Lamont, 2019). All of them are approved drugs, which should facilitate the introduction of such combinations in clinical practices.

We have shown recently that antimicrobial peptides, including pexiganan, can induce priming in bacteria, an enhanced response to the peptides when bacteria are pre-exposed to low concentrations. The consequence of priming is not only survival but an increase in tolerance and persistence (Rodríguez-Rojas et al., 2019). The use of antimetabolites could potentially abolish this property in therapeutic usage.

## Conclusions

The analysis of the pexiganan stress response by *S. aureus* has shown a global response involving several proteins known for their role in the development of resistance against antimicrobial peptides and other immune system effectors. Pexiganan has also shown a synergistic increase of antibacterial activity when it is combined with nucleoside antimetabolites. Taken together, our results suggest that pexiganan renders *S. aureus* susceptible to purine and pyrimidine analogues, which are traditionally used for cancer treatment. These antimetabolite analogues can enhance the bactericidal activity of pexiganan against *S. aureus* under the tested conditions. The significant potentiation of the pexiganan bactericidal activity and the decrease of minimal inhibitory concentrations when compared with pexiganan alone could be the basis for new formulations of pexiganan. These results are probably extendable to other antimicrobial peptides and other bacterial pathogens. Thus, the leakage of nucleotides and intermediate small metabolites or cofactors caused by cationic peptides and nucleotide metabolic pathways are common traits of bacteria-peptide interactions as proposed for the symbiont–host interface (Mergaert et al., 2017). Our results also show that understanding how antimicrobials operate and how pathogens respond to them is important to guide the design of new effective therapies. Physiological response by bacteria is informative or suggestive about additional drug combinations that can limit the chances of pathogens to evolve resistance while increasing pathogen clearance and decrease toxicity. This approach should be exploited to rationally design new antimicrobial combinations.

## Data availability

All data necessary obtained during this study are represented fully within the article. Raw data are available upon request.

## Authors’ contributions

A.R.R. and J.R. conceived the study; A.R.R., A.N., B.E.S, G.S. and B.K. performed the experiments and collected the data; A.R.R., G.S., J.J.K., B.K. and C.W. analysed the data; A.R.R. and J.R. wrote the manuscript and revised the final document. All authors agree to be held accountable for the content therein and approved the final version.

## Acknowledgements

We would like to thank Dr Dan Roizman from Freie Universität Berlin for help with Resazurin assay and critical reading of the manuscript. We would like to also thank Dr Michael A. Zasloff from Georgetown University for kindly providing the pexiganan.

---

**Figure S1.**
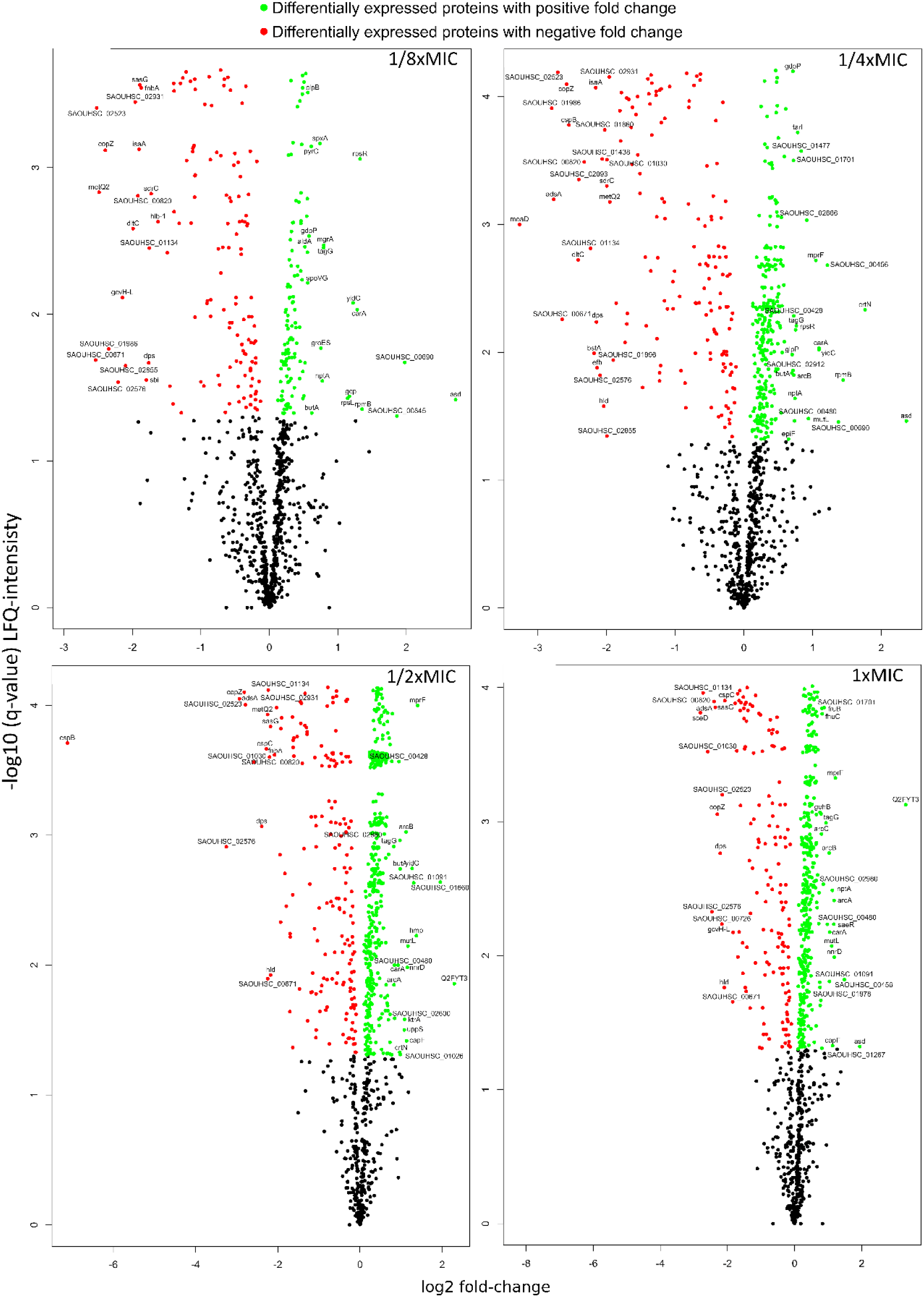
Volcano plots –log q values vs. log2 fold change of protein intensity measured by LC-MS of pexiganan treated cells with different fractions of the MIC, each compared versus untreated control). Black dots represent not significant expressed proteins while green dots show the upregulated portions and red ones represent the down-regulated fraction.

**Figure S2.**
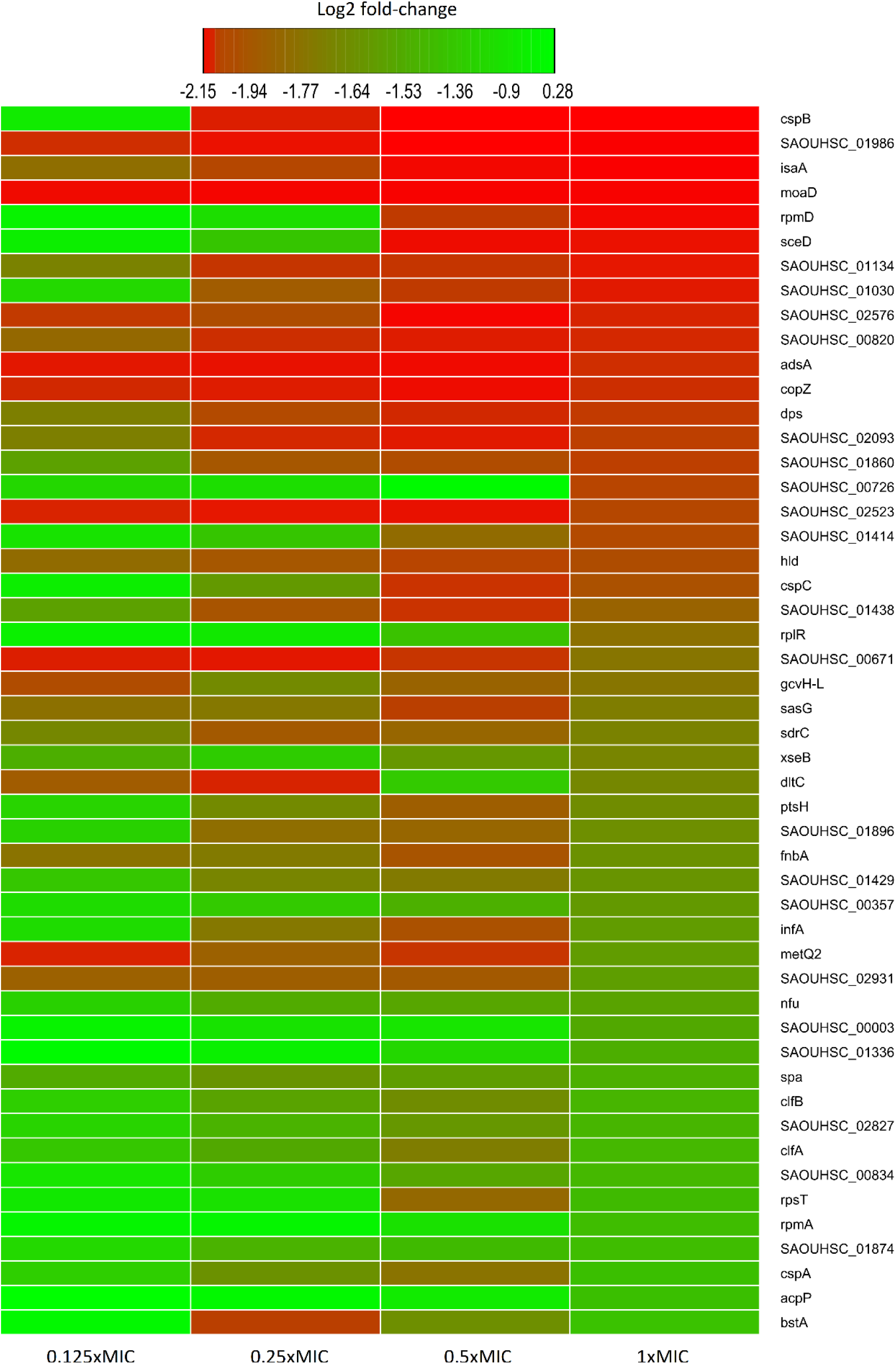
Heatmap of relative protein expression based on label-free quantification by liquid chromatography/mass spectrometry (LC-MS). Only the 50 most statistically significant down-regulated proteins are shown, taking as a reference the ones from 1xMIC pexiganan concentration (0.125, 0.25, 0.5 and 1 fractions of the minimal inhibitory concentration). Intensity ranges of the log2 fold-changes are given from highest intensity (green) to lowest (red) sorted by their values for the 1x MIC.

**Figure S3.**
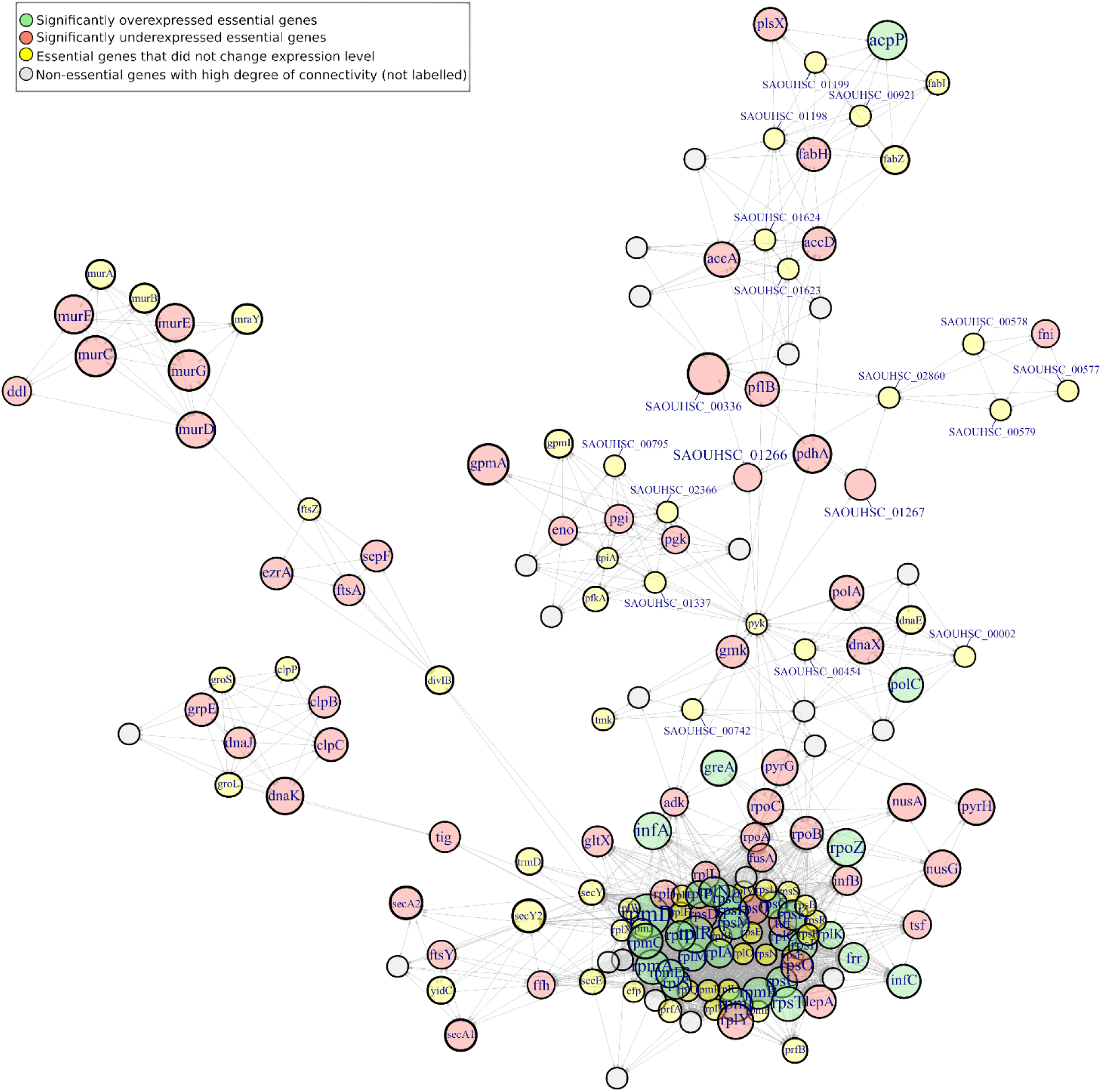
Network analysis of pexiganan stress (1xMIC) on essential genes interactome of *S. aureus* SH1000. Pale green nodes indicate upregulated proteins while pale red ones represent down-regulated ones. Grey nodes correspond with genes with a high degree of connectivity with this essential network, but they were not labelled. Note the higher proportion of downregulated genes among essential proteome while the majority of unregulated proteins are ribosomal components and thus they aggregate due to physical interaction. The interaction among nodes shows the proteome-wide impact of pexiganan stress at an inhibitory concentration.

**Figure S4.**
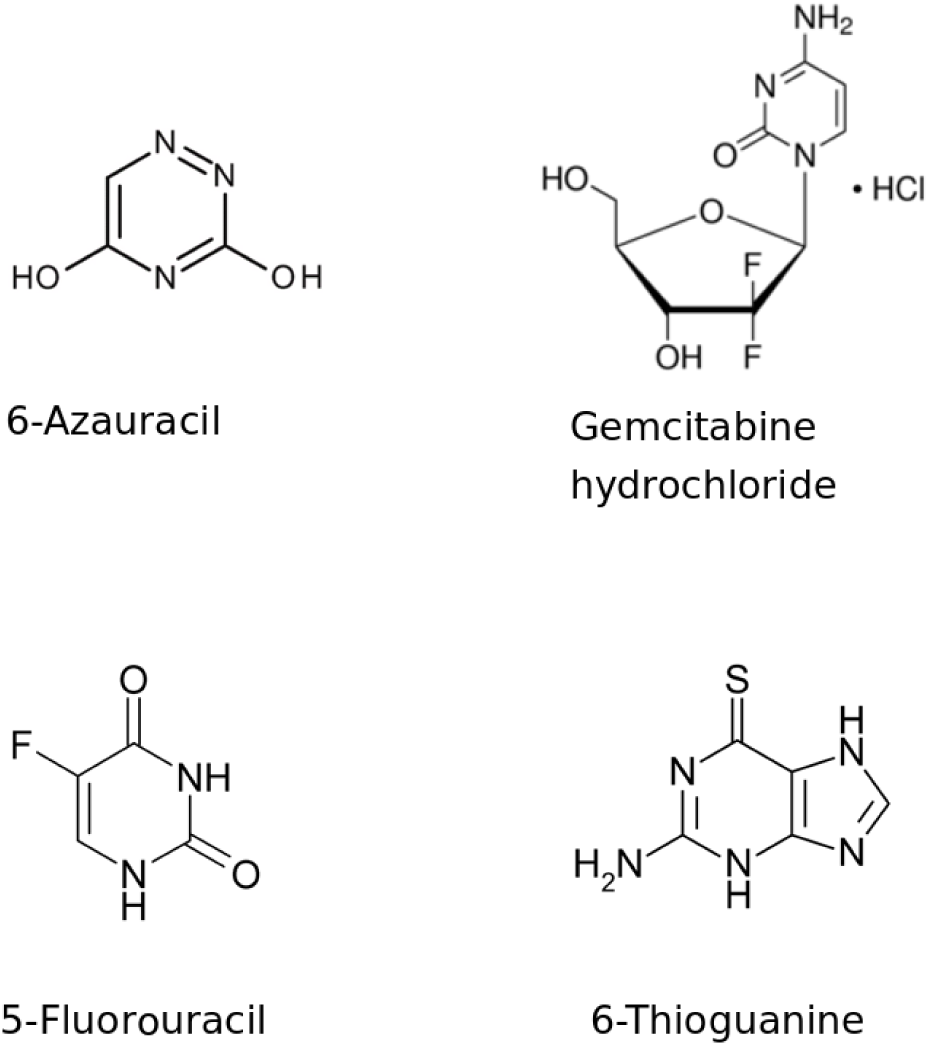
Chemical structure of nucleoside antimetabolites used in this work. Images were obtained from Wikipedia.

**Figure S5.**
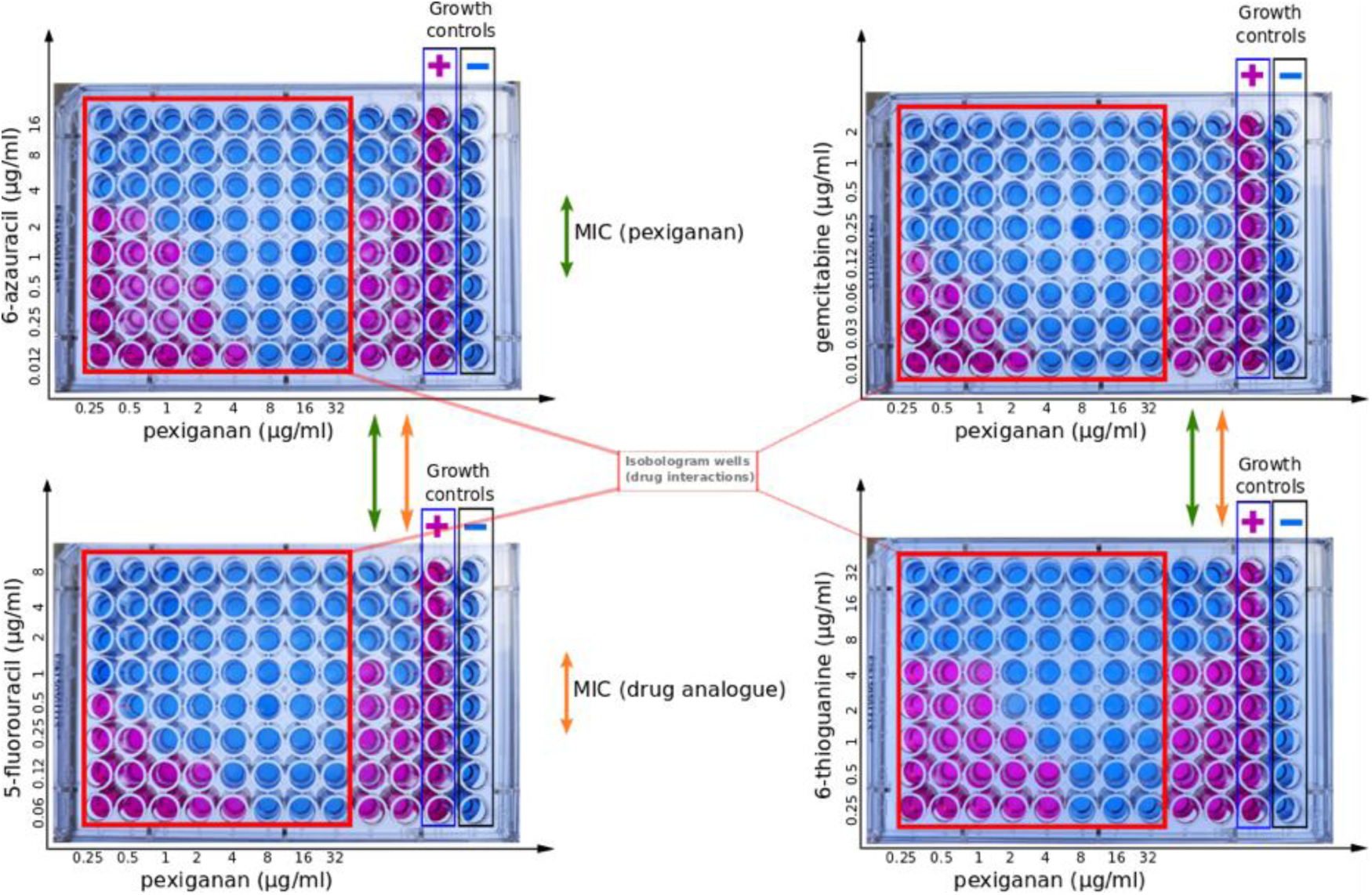
Isobologram showing the synergistic activity of pexiganan and different nucleotide antimetabolite combination against *S. aureus* SH1000. Columns with no colour change (blue resazurin) indicate no viable bacteria while colour change to purple (reduced resazurin) was considered as a sign bacterial growth.

**Table S1.**
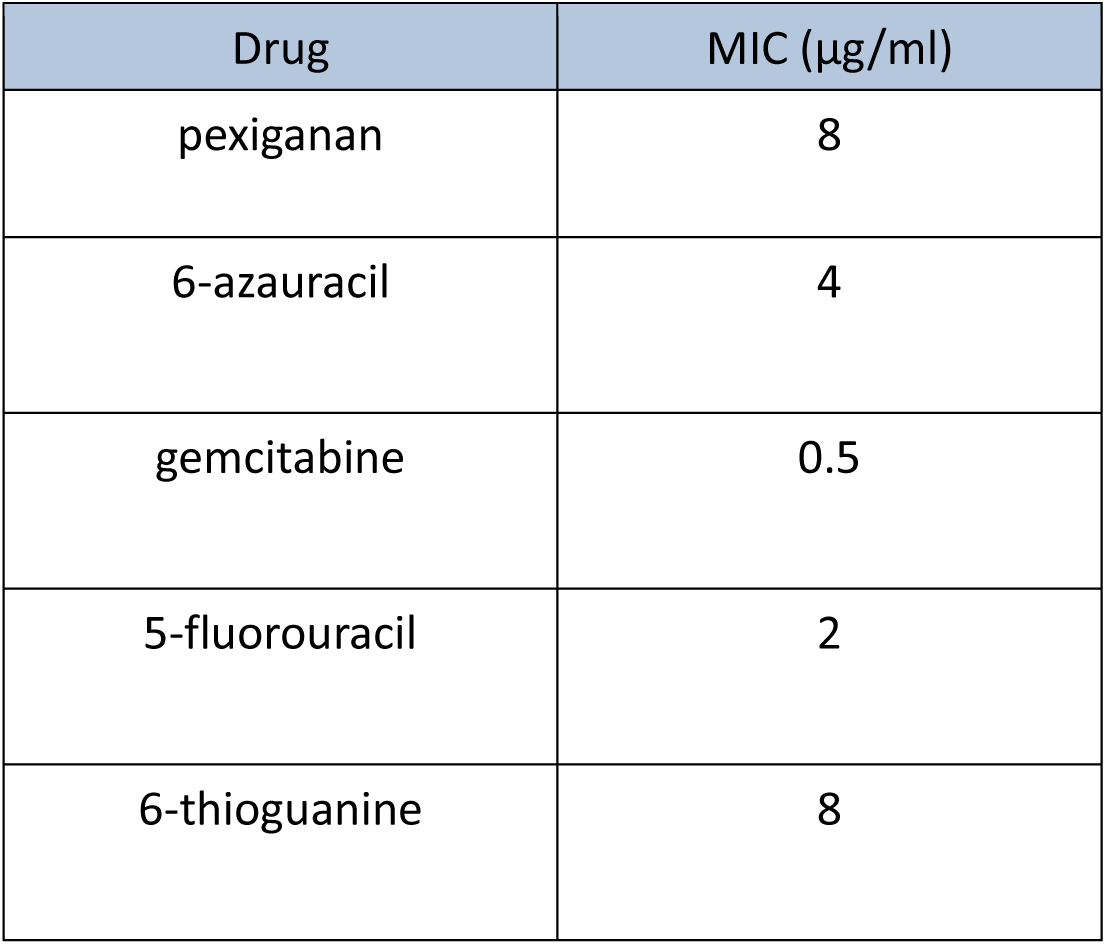
Minimum inhibitory concentration (MIC) data for *S. aureus* SH1000 to pexiganan and antimetabolite analogues used in this study.

Table S2. Output table of the proteomic experiment reporting *S. aureus* SH1000 response to pexiganan addition (0.125, 0.25, 0.5 and 1x MIC). Bacteria were sampled after 30 minutes of the treatment. Each treatment group consisted of six independent replicates and bacteria before treatment (T0) were used as control. Statistical analysis used student t-test and false discovery rate for correction of the p-values (data analysis using Maxquant (Tyanova et al., 2016a) and Perseus software (Tyanova et al., 2016b) for label-free quantification of proteins with LC-MS).

**Table S3.** Fractional inhibitory concentration (FIC) data for *S. aureus* SH1000 to Pexiganan and antimetabolite analogues used in this study.

| Drug Combination | FIC Index | Interaction type |
| --- | --- | --- |
| Pexiganan + gemcitabine | 0.051 | Synergistic |
| Pexiganan + 6- azauracil | 0.25 | Synergistic |
| Pexiganan + 5-fluorouracil | 0.037 | Synergistic |
| Pexiganan + 6-thioguanine | 0.375 | Synergistic |

